# Brain cooling marginally increases maximum thermal tolerance in Atlantic cod

**DOI:** 10.1101/658062

**Authors:** Fredrik Jutfelt, Dominique G. Roche, Timothy D Clark, Tommy Norin, Sandra A. Binning, Ben Speers-Roesch, Mirjam Amcoff, Rachael Morgan, Anna H Andreassen, Josefin Sundin

## Abstract

The physiological mechanisms determining thermal limits in fishes are debated but remain elusive. It has been hypothesised that loss of motor function observed as a loss of equilibrium during an acute thermal challenge is due to direct thermal effects on brain neuronal function. To test this hypothesis, we mounted cooling plates on the head of Atlantic cod *(Gadus morhua)* and quantified whether local cooling of the brain increased whole-organism critical thermal maxima (CT_max_). Brain cooling reduced brain temperature by 2–6°C and increased CT_max_ by 0.5–0.7°C relative to instrumented and uninstrumented controls, suggesting that direct thermal effects on brain neurons might contribute to setting upper thermal limits in fish. However, the improvement in CT_max_ with brain cooling was small relative to the difference in brain temperature, demonstrating that other mechanisms (e.g., failure of spinal and peripheral neurons, or muscle) may also contribute to controlling acute thermal tolerance in fishes.

**Summary statement:** We tested whether brain temperature sets the upper thermal limit in a fish. Selectively cooling the brain during whole-organism thermal ramping marginally increased thermal tolerance.

## INTRODUCTION

Warming from climate change is increasing mean temperatures as well as the frequency and severity of heat waves (Seneviratne et al., 2014). Severe heat waves can lead to mass mortality in aquatic ecosystems, (Wegner et al., 2008), and may thus constitute a strong selection force (Sunday et al., 2014), potentially even in thriving populations (Sandblom et al., 2016). The vast majority of aquatic ectothermic water-breathers have the same body temperature as the surrounding water. With heat waves on the rise in many aquatic systems, thermal challenges are likely becoming an increasingly important selection force for fishes (Seneviratne et al. 2014).

Despite more than a century of research on acute thermal challenges in fishes, the precise mechanisms that lead to loss of equilibrium (LOE) remain elusive (Beitinger & Lutterschmidt, 2011; Carter, 1887; Davy, 1862). In an experiment by Friedlander et al. (1976), goldfish *(Carassius auratus)* showed the same critical thermal minimum (CT_min_), critical thermal maximum (CT_max_), and behavioural responses to temperature when only the brain temperature was manipulated (by the use of thermodes mounted on top of the cerebellum) as when the ambient water temperature was manipulated (Friedlander et al., 1976). The study by Friedlander et al. suggests that the effect of temperature on neural function may be responsible for LOE during acute warming. However, this idea remains largely unexplored. To test whether brain temperature is the main controller of LOE at the acute upper thermal limit, we mounted custom-made cooling plates on the skin above the brain of Atlantic cod *(Gadus morhua).* The plates were flushed with either ambient temperature water or chilled water while the fish underwent a thermal ramping protocol. We predicted that fish with cooled brains would show LOE at higher water temperatures than fish with brains maintained at the ambient water temperature.

## MATERIALS AND METHODS

### Experimental animals

Juvenile Atlantic cod were cage-caught in the waters off Lysekil, Sweden, in June 2017 and brought by boat to the Sven Lovén Centre for Marine Infrastructure, Kristineberg, University of Gothenburg, Sweden. At the Centre, the fish were kept in two 1000 L tanks with flow-through seawater pumped from 30 meters depth. The thermoregulated water was increased from 10.7°C – the natural ambient temperature at time of capture – to the target acclimation temperature of 14°C over a period of three days. The fish were then acclimated to 14°C for three weeks before the experiments commenced (actual mean ± SD temperatures were 13.74 ± 0.97°C in holding tank one and 13.76 ± 0.98°C in holding tank two). The cod were fed blue mussels *(Mytilus edulis)* and shrimp *(Pandalus borealis)* every second day. Artificial plastic plants and cut PVC pipes were provided in the tanks for shelter. The light cycle was set to L 18 h: D 6 h, following natural conditions. The experiments were conducted in accordance with ethical permit Dnr103-2014, from the Swedish Board of Agriculture.

### Brain coolers

Custom-built brain coolers (Fig. 1A) were machined out of a solid block of aluminium using a CNC mill by the workshop at the Norwegian University of Science and Technology, Trondheim, Norway. The vertical and horizontal holes for the U-shaped pipe loop running through each brain cooler were drilled, and the horizontal hole was plugged at each end to form the loop. Two different sizes of brain coolers (15 × 6 mm and 20 × 10 mm) were used to accommodate the range of fish sizes used in the experiment (Fig. S2). The coolers were attached to the top of the head of the cod using cyanoacrylate glue and silk sutures (Fig. 1B), and connected to a thin flexible silicone tubing (2 mm ID, 4 mm OD) that allowed water to be flushed through the coolers to control their temperature (Fig. 1C).

**Fig. 1.**
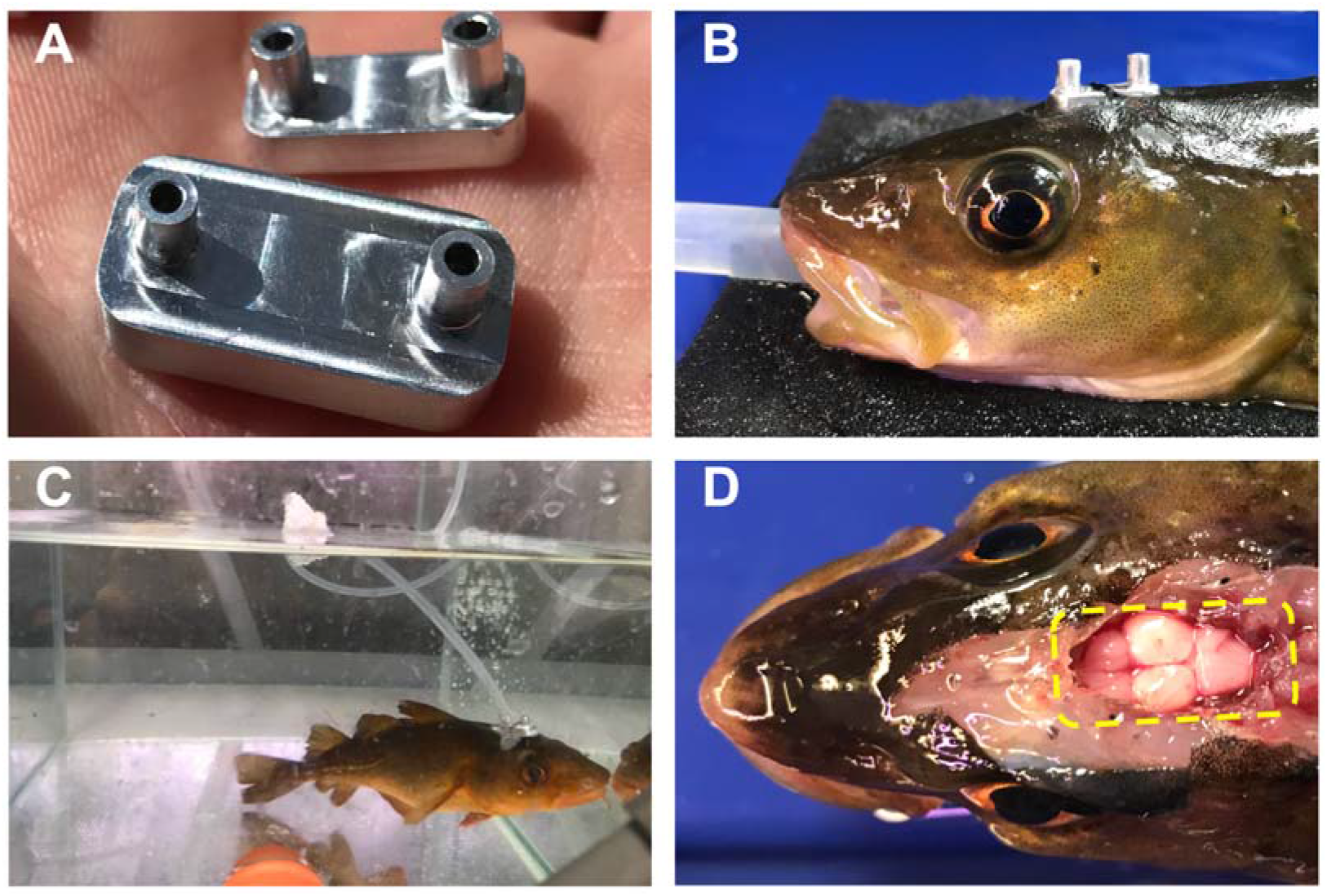
Design and attachment method of the brain coolers. (A) Solid aluminium brain coolers with a u-shaped loop running through the block, allowing for water flow through. (B) Brain cooler mounted on the dorsal cranium of an Atlantic cod, using cyanoacrylate glue and sutures. (C) A thin and flexible silicone tubing was used to flush the brain cooler with ambient or cold water while allowing normal behaviour during a CT_max_ challenge. (D) The top of a euthanised cod with the cranium opened, showing the cooled brain regions (the yellow rectangle indicates the position of the cooler).

To attach the brain coolers, fish were anesthetised in a tank using MS-222 (50-60 mg L^-1^) and then placed on a surgery bench where the gills were ventilated via silicone tubing (Fig. 1B) with recirculated water with a maintenance dose of MS-222 (30 mg L^-1^). After carefully rinsing and drying the attachment area on top of the head to remove mucous, a brain cooler was attached to the skin (Fig. 1B). This assured close connection between the brain cooler and the head of the fish, allowing efficient heat transfer from the head to the cooler. Fig. 1D shows the position of the cooler relative to the brain.

### Brain cooling validation

In addition to the experimental fish, three fish (total length = 24.1 ± 2.7 cm, body mass = 122.2 ± 52.8 g; means ± SDs) were used to test the cooling capacity of the brain coolers on brain tissue. These fish were terminally anesthetised and instrumented with thermocouples (TC-08, Picotech, Cambridgeshire, UK) in different parts of the brain (different points in different fish) and subsequently thermally ramped (Fig. 2). Close to the cranium, the cooling effect was 6°C, while the ventral side of the brain was cooled by as little as 2°C.

**Fig. 2.**
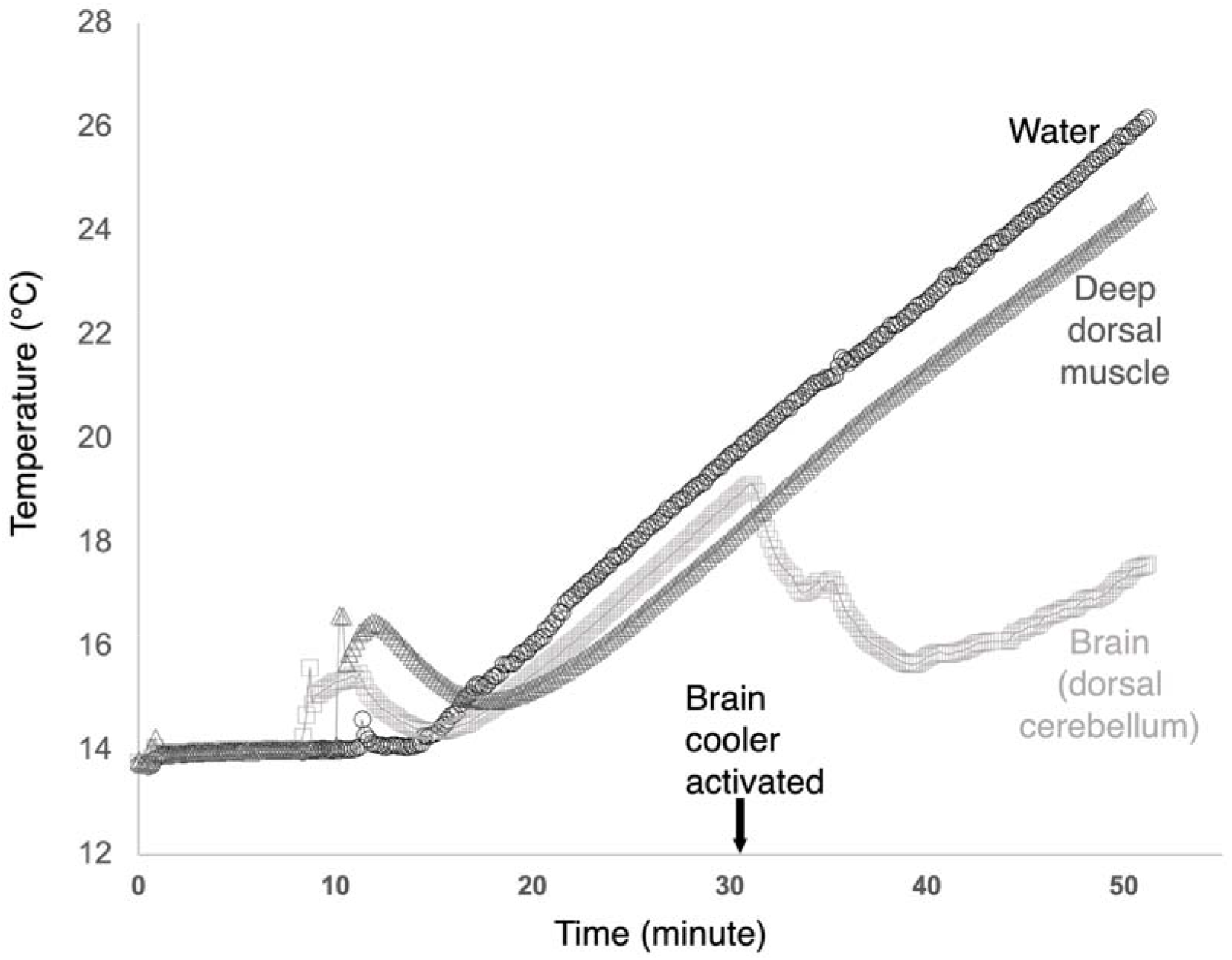
Brain cooling validation. A raw trace example of three thermocouples during a thermal ramping CT_max_ protocol in a pilot experiment fish. One thermocouple was placed in the aquarium, showing the ambient water temperature (black circles). Another thermocouple was placed inside the deep dorsal muscle of a terminally anesthetised Atlantic cod in the aquarium during thermal ramping (dark grey triangles). The third thermocouple was placed adjacent to the cerebellum of the same fish (light grey squares).

### CT_max_ setup

CT_max_ experimentation methodology has been thoroughly described and validated previously (Morgan et al., 2018), and is briefly described below. Four aquaria (30 × 30 × 25 cm, two-thirds filled) were used in parallel for testing the acute maximum thermal tolerance of the cod. The aquaria each had an overflow connected to a heating sump in which water temperature was ramped using a 500 W titanium heater (Aquamedic, Bissendorf, Germany). A large water pump (DC runner 9.1, Aquamedic, Bissendorf, Germany) with the flow split four-ways supplied each of the four aquaria with 3.75 L min^-1^ of recirculating water. The heating sump had heavy aeration to ensure gas equilibrium with the atmosphere. The temperature in the aquaria was continuously recorded by thermocouple loggers (TC-08, Picotech, Cambridgeshire, UK) connected to a PC.

The thermal ramping rate during the CT_max_ experiments was 10°C h^-1^. The brain coolers of the cooling treatment group were supplied with ice-cold seawater, pumped from an adjacent container (Eheim Universal 1046 pump, Eheim GmbH, Germany). The brain coolers of the ambient temperature treatment group (i.e., instrumented control group) were supplied with ambient ramping-temperature seawater pumped from a control aquarium (Eheim Universal 1046 pump, Eheim GmbH, Germany). To avoid cold shock to the brains of the cooling treatment group at the start of thermal ramping, the pumps to the coolers were only activated once ambient water temperature had increased by 3-4°C. The CT_max_ test of the control treatment group followed the same methods with the exception that they were not instrumented with brain coolers. The sample size, total length, and body mass of cod from the three treatment groups are presented in Table 1.

**Table 1.** Critical thermal maximum in °C with and without two statistical outliers (CT_max_, CT_max_NO_, respectively), total length (cm), and body mass (g) of Atlantic cod in three groups: control (uninstrumented fish, n=18), ambient (instrumented control group: fish mounted with brain coolers receiving ambient ramping-temperature water, n=9), and cooled (treatment group: fish mounted with brain coolers receiving cooled water, n=11). The mean and standard deviation (SD) are shown for each group, as well as the mean difference (Δ) between groups and the 95% bootstrapped confidence interval (CI).

| | **control**<br>(mean ± SD) | **ambient**<br>(mean ± SD) | **cooled**<br>(mean ± SD) | **ambient - control**<br>($\Delta$ [95% CI]) | **cooled - control**<br>($\Delta$ [95% CI]) | **cooled - ambient**<br>($\Delta$ [95% CI]) |
| --- | --- | --- | --- | --- | --- | --- |
| **$CT_{max}$** | 25.68 ± 0.80 | 25.82 ± 0.54 | 26.33 ± 0.49 | 0.14 [-0.31–0.67] | 0.64 [0.25–1.18] | 0.51 [0.08–0.95] |
| **$CT_{max\_NO}$** | 25.82 ± 0.58 | 25.96 ± 0.36 | 26.33 ± 0.49 | 0.15 [-0.20–0.51] | 0.51 [0.12–0.89] | 0.37 [-0.01–0.71] |
| **Total length** | 21.98 ± 3.24 | 24.26 ± 3.04 | 22.95 ± 2.31 | 2.27 [-0.22–4.45] | 0.97 [-1.17–2.76] | -1.30 [-3.64–1.02] |
| **Body mass** | 94.90 ± 45.47 | 120.53 ± 39.82 | 110.07 ± 38.78 | 26.5 [-8.80–54.60] | 15.20 [-17.30–42.30] | -10.50 [-43.60–22.60] |

The fish were closely monitored for behavioural changes during thermal ramping. Some individuals regurgitated food during ramping. Fish were deemed to have reached their CT_max_ at the temperature where they exhibited LOE and were unable to right themselves within three seconds (Morgan et al., 2018). At this point, the time, temperature, and fish mass were recorded, and the fish was immediately killed by a blow to the head.

### Statistical analyses

To avoid common pitfalls of p-values (Halsey et al., 2015), we examined differences in fish size and CT_max_ among groups using estimation statistics rather than null hypothesis tests (Ho et al., 2018; Halsey, 2019). We present all data points, group means and standard deviations, and treatment effect sizes with 95% confidence intervals computed from 5,000 bootstrapped samples. Statistics and plots were produced using the ‘dabestr’ package (Ho et al., 2018) in R v3.5.0 (R Core Team, 2018). Two statistical outliers were removed from the dataset to examine their influence on statistical outputs (Fig. S1). The data and analysis script are publicly available on the repository figshare (https://figshare.com/s/13ea251dc8c883e0d775) and were made available to the editors and reviewers upon submission.

## RESULTS AND DISCUSSION

The brain coolers successfully reduced brain temperature despite being attached to the skin, on the outside of the skull. The thermocouples, placed at different locations around the dorsal cranium, recorded temperature reductions of 2–6°C depending on their distance from the brain cooler (Fig. 2). Brain cooling did not appear to affect whole body temperature during thermal ramping, suggesting that the cooling was localised and that the temperature difference between the brain and deep muscle was maintained throughout the thermal ramping (Fig. 2). This demonstrates that the external brain coolers functioned as intended. External brain coolers are, therefore, effective and practical tools for investigating effects of brain temperature on fish physiology and behaviour in a less invasive way than previous methods using thermodes implanted inside the cranium (Friedlander et al. 1976).

There was no statistical difference in body length and mass among cod in our three experimental groups: fish without brain coolers (control group), fish with brain coolers flushed with ambient ramping-temperature water (instrumented control group) and fish with brain coolers flushed with cool water (treatment group) (Table 1). Cod in the treatment group tolerated higher temperatures before reaching LOE than cod in the control group (mean difference in CT_max_ of 0.64°C, 95% CI = 0.25–1.18°C) and cod in the instrumented control group (mean difference in CT_max_ of 0.51°C, 95% CI = 0.08–0.95°C) (Table 1, Fig. 3). The small difference in CT_max_ between the control and instrumented control groups (0.14°C, 95% CI = −0.31–0.67°C) suggests that the instrumentation procedure had a minimal effect on LOE. Removing a statistical outlier in the control group (23.4°C) and one in the instrumented control group (24.7°C) reduced the mean difference in CT_max_ with the treatment group to 0.51°C (95% CI = 0.12–0.89°C) and 0.37°C (95% CI = −0.01–0.71°C), respectively (Table 1, Fig. S1).

**Fig. 3.**
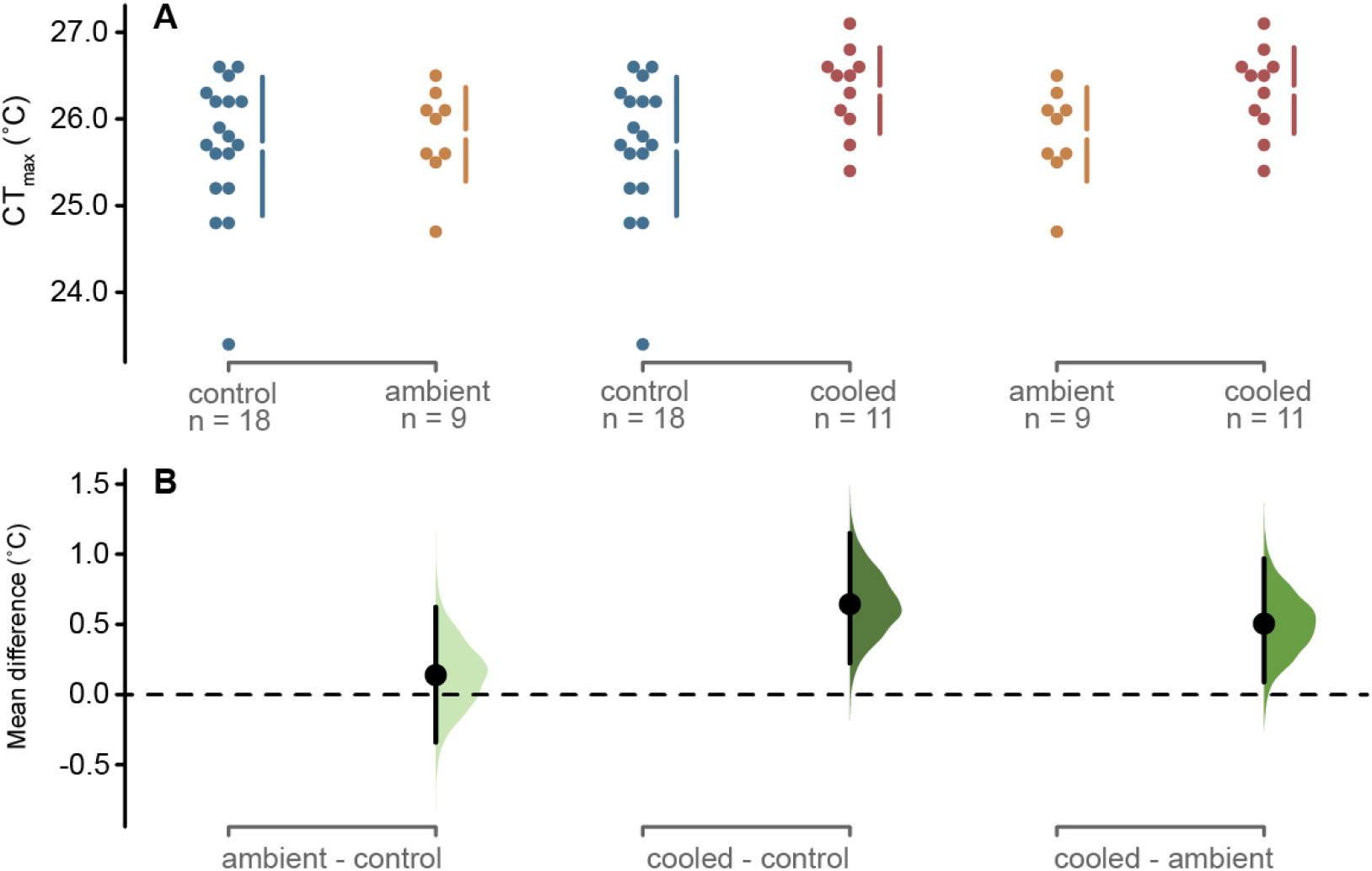
Critical thermal maximum (CT_m_ax) measured as loss of equilibrium temperature in three groups of Atlantic cod. (A) CT_max_ values of the uninstrumented group (control) are shown in blue, the instrumented control group (ambient) in orange (fish were mounted with brain coolers receiving ambient, ramping-temperature water), and the treatment group (cooled) in red (fish were mounted with brain coolers receiving cooled water). Vertical bars indicate the standard deviation around the group mean (shown as a gap). (B) Cumming estimation plots (Ho et al., 2018) showing the mean differences in CT_max_ among the three groups (i.e., effect sizes; black dots), the distribution of these effect sizes obtained through nonparametric bootstrap resampling (5,000 samples), and their 95% confidence intervals (black bars).

The elevated CT_max_ in brain cooled fish supports our prediction that cooling the brain increases whole-organism thermal tolerance. Our results are also in accordance with an earlier study in which manipulation of brain temperature in goldfish caused the same behavioural effects and LOE temperatures as did warming the whole animal (Friedlander et al., 1976). These results suggest that the brain could be an important organ affecting thermal limitation during acute thermal challenges in fish. However, the cooling effect of the brain coolers in our study was large (2–6°C depending on the brain region), while the increase in CT_max_ was comparatively small (0.5–0.7°C). We would have expected a larger increase in whole-organism CT_max_ if the brain was the sole organ controlling LOE. As CT_max_ was only marginally elevated by brain cooling, it is possible that peripheral neurons and muscles could potentially have very similar thermal limits as the brain. One approach to disentangling variation in thermal tolerance between these different organs and cell types could be selective cooling, using externally mounted coolers similar to those used here, or by implanting thermodes for cooling specific tissues (e.g. brain, muscle, heart) (Friedlander et al., 1976). Another path could be *in situ* or *in vitro* characterisation of thermal limits in partitioned organ systems (Ern et al., 2015).

During acute thermal ramping, fish can show increasing spontaneous movements at higher temperatures, before ceasing righting movements at LOE (Beitinger and Lutterschmidt, 2011). As the cod in this study approached LOE, they suddenly appeared to reduce fin movements (unquantified personal observation), which led to a loss of righting behaviour. This reduction in fin movements indicated loss of motor control, which could be caused by muscle dysfunction, neuronal dysfunction, or both simultaneously. If the direct effect of high temperature on skeletal muscle contractility was the cause of LOE, then we should not have been able to affect CT_max_ with the brain coolers. Conversely, if the brain is solely responsible for setting thermal limits, we would have observed a larger effect of brain cooling on CT_max_. Thus., the most parsimonious explanation for our observations seems to be that the central and peripheral nervous systems, and potentially the muscle, have very similar thermal limits.

The ‘oxygen- and capacity-limited thermal tolerance’ (OCLTT) hypothesis suggests that upper thermal limits are set by the inability of ectothermic organisms to deliver a sufficient supply of oxygen to the tissues. When warming pushes an animal’s metabolic rate to levels where oxygen delivery is insufficient, tissue hypoxia ensues (Pörtner and Knust, 2007). The OCLTT hypothesis remains controversial, yet can be used to form testable predictions (Clark et al., 2013; Jutfelt et al., 2018). Accordingly, OCLTT predicts that brain hypoxia would cause LOE during heat challenges. In fish, heart failure during thermal ramping (Ekström et al., 2016) due to cardiac muscle hypoxia has also been suggested to contribute to upper thermal limits (Farrell, 2009). Collapsing circulation would consequently lead to brain or muscle hypoxia that causes LOE. As Atlantic cod in the present experiment did not show a major increase in CT_max_ with brain cooling, our results do not refute OCLTT predictions. However, as the cooling was local to the brain, cooling should not have protected against cardiac collapse (Farrell, 2009). The slight increase in CT_max_ due to brain cooling thus suggests that a direct thermal effect on neuronal function is a candidate mechanism involved in setting acute thermal limits in fish.

## Acknowledgements

We thank Bengt Lundve and the Royson family for fish collections, and the staff of the Sven Lovén Centre for Marine Infrastructure, Gothenburg University, Kristineberg, for technical assistance. We thank Elisabeth Valberg at the NTNU mechanical workshop for brain cooler blueprints and construction.

## Author contributions

FJ designed and performed the experiment with input from all authors. JS, TN, MA, and BSR cared for the fish. DGR and JS analysed the data. FJ wrote the manuscript draft with significant contributions and final approval from all authors.

